# Control of cartilage homeostasis and osteoarthritis progression by mTORC1/4E-BP/eIF4E axis

**DOI:** 10.1101/768457

**Authors:** Olga Katsara, Mukundan Attur, Victoria Kolupaeva

## Abstract

As world population growing older, burden of age-related conditions soars. One of them is osteoarthritis (OA), a debilitating joint disease with no effective treatment. Articular cartilage degeneration is a central event in OA, and changes in expression of many genes in OA cartilage are well-documented. Still, the specific mechanisms are rarely known. We showed that in OA cartilage the increased abundance of many proteins, including extracellular matrix protein Fibronectin (Fn1) and an orphan nuclear receptor Nr4a1 is translationally controlled and requires inactivation of 4EBP, an inhibitor of cap-dependent translation. Importantly, intra-articular injection of the translation inhibitor 4E1RCat reduces Fn1 and Nr4a upregulation in a rodent OA model and delays cartilage degeneration. Our results support the hypothesis that maintaining proper translational control is an important homeostatic mechanism, the loss of which contributes to OA development.

## INTRODUCTION

Extracellular matrix (ECM) determines many unique mechanical properties of articular cartilage. Fibronectin (Fn), an ECM protein, is a minor component of healthy cartilage; but in OA cartilage, there is an up to tenfold accumulation of the protein (1, 2). Fn fragments induce pro-inflammatory cytokines, matrix-degrading metalloproteinases (MMPs), and suppress collagen and aggrecan expression leading to the loss of the biomechanical properties of cartilage (3–6). Both transcriptional (7) and post-transcriptional (2) regulation of Fn expression was proposed in different biological contexts, yet, the mechanism leading to an increase in Fn abundance in OA cartilage is not known. Different groups reported conflicting data on the Fn mRNA expression in OA cartilage. . Few groups reported a modest increase in Fn RNA expression in OA cartilage compared to normal tissue (8, 9), but the majority found no significant difference (10–14). We, therefore, investigated if translational control is responsible for Fn1 abundance in OA cartilage.

The importance of translational control is emerging in understanding the vast majority of biological processes, particular as the assessment of mRNAs expression level often does not completely correlate with proteomic data. It is becoming apparent that in many cases the transcriptome does not accurately represent the proteome (15). Recent publications began to address the role of post-transcriptional control of gene expression in chondrocyte homeostasis and OA progression (16–19). We used IL-1β stimulated primary articular chondrocytes as they possess many characteristics of OA chondrocytes. Importantly, IL-1β production is associated with cartilage degeneration in OA (20), and it was shown that inflammatory molecules produced by synovium can induce ECM loss (21). The importance of IL-1 signaling was also highlighted in animal models of posttraumatic OA (22–24). Potential clinical relevance of identified translationally controlled targets, such as Fn1 and an orphan receptor Nr4a1, was tested in diseased cartilage and in a rat model of post-traumatic OA. To demonstrate the importance of translational control for cartilage homeostasis, we injected intra-articularly an inhibitor of cap-dependent translation. This treatment reduced Fn1 and Nr4a1 upregulation and significantly delayed OA progression. Our findings demonstrate that translational control, acting at Fn and Nr4a1 among other targets, is essential for maintaining cartilage homeostasis.

Recent publications from two independent groups have highlighted the importance of IL-1 signaling in various small and larger animal models of posttraumatic OA (22–25). For example, IL-1 receptor antagonist therapy significantly improved cartilage, synovial membrane parameters, and disease-modifying effects in the equine OA model (26). The role of translation control in the inflammatory response was examined in macrophages (27), but never in chondrocytes. We therefore used polysome profiling to assess the translational landscape of IL-1β treated primary articular chondrocytes. In polysomal profiling, actively translated mRNAs bound by several ribosomes (polysomes) are separated from “free” mRNA and the 80S monosomes by sucrose gradient centrifugation. This allows determining the translational status of a specific mRNA. Potential clinical relevance of several identified translationally controlled targets, such as Fn and an orphan receptor Nr4a1, was tested in diseased cartilage and in a rodent model of post-traumatic OA. To demonstrate the importance of translational control for cartilage homeostasis, we injected intra-articularly an inhibitor of cap-dependent translation. This treatment significantly delays OA progression. Our findings therefore reveal that translational control, acting at Fn and Nr4a1 among other targets, is essential for maintaining cartilage homeostasis. Moreover, targeting translational apparatus represents a new strategy to treat/prevent OA, particularly as it opens up the possibility to target simultaneously a pool of functionally distinct proteins.

## MATERIALS AND METHODS

### Isolation and characterization of translationally active pool of mRNAs by polysome fraction analysis, library preparation, and sequencing

RAC was isolated and treated at the first passage as described previously (18). For 10 final minutes, cells were incubated with Cyclohexamide (100ug/ml) at 37°C. Cells were collected, washed twice with ice-cold PBS supplemented with Cyclohexamide (100ug/ml) and lysed on ice for 10min in buffer “A” (SI). 2OD units of extract (3 independent experiments combined) were layered over 5-50% linear sucrose gradients in buffer “B” (SI) and centrifuged at 38,000 rpm for 2h at 4°C. Gradients were fractionated while monitoring RNA absorbance at 254nM. RNA was isolated from pooled polysome fractions by phenol-chloroform extraction (see SI for details). Library preparation, sequencing, and downstream analysis were done as described in SI.

*qPCR, Immunohistochemical and IF analyses* were done as described in SI, using antibodies listed in Sup.Table 10.

### Construction of 5’UTR-Luc reporter plasmids and reporter assay

5’UTRs of Mmp13, Aspn, Sulf1, Nr4a1 and Fn1 were amplified from total mRNA using specific primers (Sup.Table 9). Same primers with extra nts containing Xho1 (5’end) and Kpn1 (3’end) sites were used for the second round of PCR and following digestion, an insert was ligated into pIS0m (mutated, see SI for details) vector. 5’UTR Nr4a1 mutant was made using Q5^®^ Site-Directed Mutagenesis Kit (NEB BioLabs) according to the manufacturer protocol. pIS0m vector was made from original pIS0 vector (Addgene) by adding additional restriction sites (Xho1, EcoR1 and Kpn1) right after SV40 promoter and immediately before firefly Luc using Q5^®^ Site-Directed Mutagenesis Kit (NEB BioLabs) according to the manufacturer protocol.

### Experimental OA in a rodent model

All animal experiments were performed in accordance with the regulations of the Institutional Animal Care and Use Committee at NYU School of Medicine. Experimental OA model is described previously (18) and in SI.

### Rat chondrocyte culture

Articular cartilage was isolated and cultured as described in (18). Once the cells reached confluence, they were starved for 16h and treated with 10ng/ml of IL1-β. For the inhibitor experiments cells were pretreated with DMSO, Ku-063794 (indicated concentrations), or Nr4a1 inhibitor DIM-C-pPhOH (10μM; Tocris Bioscience) for 1h and then treated with IL1-β for 24hrs. Transfections with Nr4a1 siRNA (nt1701; Sigma) were performed using Lipofectamine RNAiMax according to the manufacturer’s protocol. MISSION siRNA Universal Negative Control was used as a control. Cells were starved for 16h in serum-free medium 48h post-transfection and then treated with IL1-β.

### Microarray analysis of human chondrocytes

Microarray analysis of the human knee cartilage was done as previously described (*28*).

### Statistical analysis

GraphPad Prism 8.03 was used for statistical analysis as indicated in figure legends.

## RESULTS

### Fn expression is controlled at the translational level in articular chondrocytes stimulated with IL-1β

Elevated Fn protein expression in OA cartilage is well-documented (1, 2, 29), we therefore first examined Fn expression at the mRNA and protein levels in human knee OA cartilage. Since it is difficult to obtain age-matched normal cartilage, we analyzed lesional and non-lesional areas of OA cartilage isolated from the same patient. The degree of cartilage degeneration reflected in loss of proteoglycans and fibrillation was visualized with Safranin O staining. In lesion areas, we observed increased immunostaining of FN1 and MMP13, which is a known player in OA pathology. Consistent with previous reports we observed significant upregulation of MMP13 but not FN1 mRNA when compared human OA (n=12) and non-OA (n=9) cartilage (Fig.1A; Sup.1A). A similar result was obtained by analyzing human Affymetrix U133A microarray data (Sup.1B). These data indicate that increased FN1 protein abundance is controlled post-transcriptionally in human OA cartilage. We observed similar results in a rat anterior cruciate ligament transection (ACLT) model of post-traumatic OA. Animals were sacrificed 20 weeks post-op. Mankin scoring system of Toluidine Blue stained tissue was used to assess the severity of cartilage lesions (with numbers ≥10 indicating the onset of OA) (30) (Sup.1C). Fn1 mRNA level was not changed in operated compared to sham joints, but IHC analysis confirmed increased Fn1 (and Mmp13) protein level in OA cartilage (Fig.1B, Sup.1D, 1E).

**Figure1.**
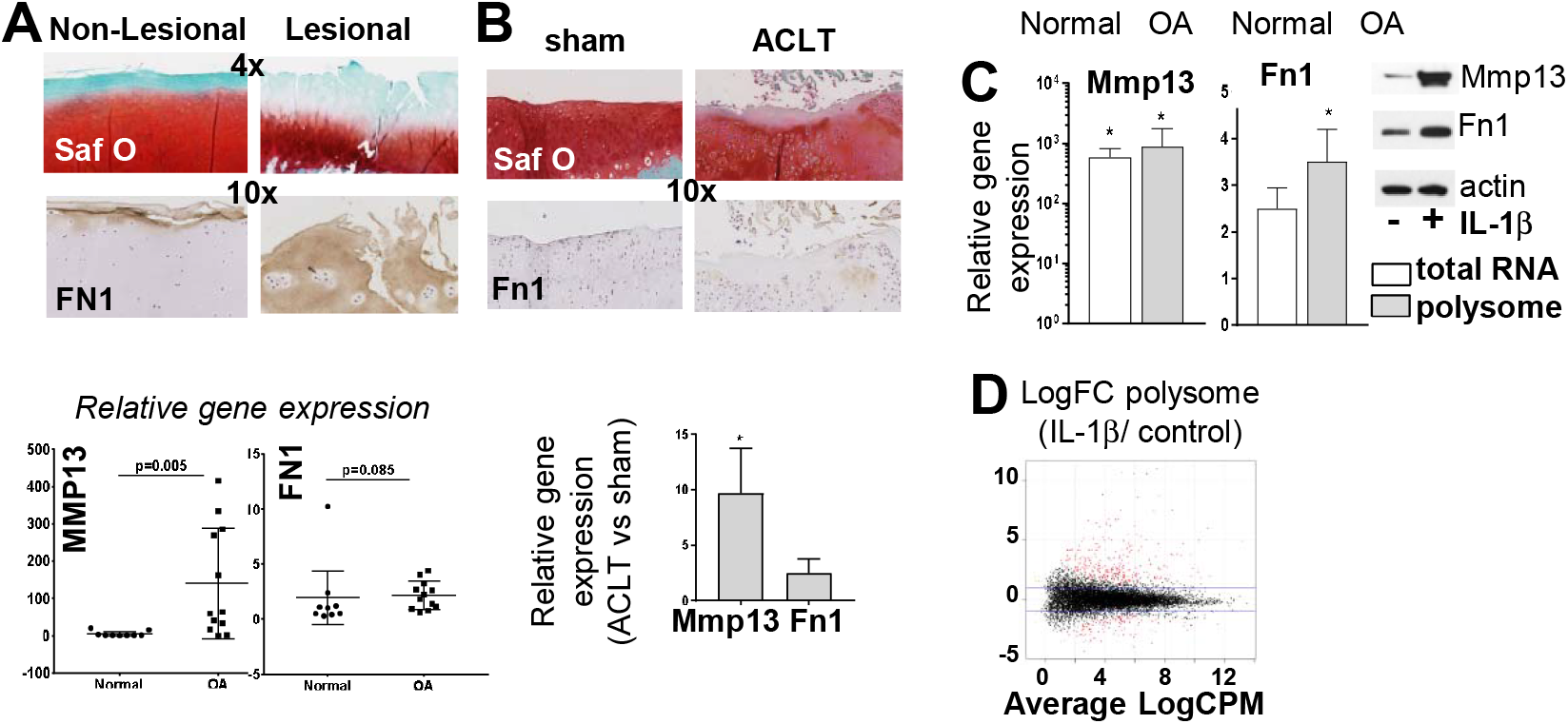
Fn1 expression in osteoarthritic (OA) cartilage. **(A)** Representative Safranin O staining of non-lesional and lesional areas of cartilage from the same patient (n=4). The lesional areas exhibit chondrocyte clustering and decreased proteoglycan staining. IHC shows accumulation of Fn1 in lesional sections. Graphs show levels of gene expression, analyzed by qPCR, as a fold change (FC) (normal (n=9) vs OA (n=12) cartilage), Mann-Whitney test. **(B)** Safranin O staining and IHC analysis of Fn1 in the rat ACLT model. Bars show levels of gene expression, analyzed by qPCR as a FC (ACLT vs sham; isolated from raw samples n=4, *t*-test,*p ≤ 0.05). **(C)** qPCR analysis of total and polysome-bound mRNA from Rat articular chondrocytes (RAC) is shown as a FC (IL-1β treated vs untreated) n=3, *t*-test,*p ≤ 0.05). Protein levels are evaluated by WB (20ug of total protein). **(D) Polysome profiling of IL1β treated RAC**. Smear plot of differential polysome loading magnitude vs. coverage (2 controls vs. 2 treated (combined from 3 independent experiments). Differentially translated transcripts are marked in red. LogFC for treated vs. control is plotted against average log expression values (standardized read counts). A two-sided t-test was performed correcting for multiple testing by controlling for FDR at 1%.

As an inflammation component is well described in OA pathology, we used IL-1β treated primary rat articular chondrocytes (RAC) as an *“in vitro”* model that displays many characteristics of OA chondrocytes, including secretion of Mmps (31). For polysome analysis untreated and IL-1β treated RAC lysates were fractionated on a sucrose gradient. Actively translated mRNAs are present in the polysome (heavy) fractions, while translationally repressed mRNAs are excluded (Sup.2A). Actively translated (bound to polysomes) mRNAs (mRNApoly) were isolated from polysome fractions, and Fn1 and Mmp13 mRNA levels were examined. IL-1β signaling considerably increases mRNApoly levels for both Mmp13 and Fn1 (Fig.1C), but only total Mmp13, not Fn1 mRNA level is significantly increased, indicating that Fn1 protein abundance is likely mediated by translation control (Fig.1C). This raises an important question; are there other translationally regulated proteins in OA-like chondrocytes? Their (not all) mRNAs would primarily benefit from accelerated protein synthesis in OA cartilage (18). To answer this question, we next characterize the translatome of IL-1β treated RAC using deep sequencing of polysome-associated mRNAs (mRNApoly).

### Polysome profiling in articular chondrocytes identifies dynamic regulation of the translatome by IL-1β

Primary RAC were treated for 24hrs with IL-1β, and polysomal fractions were sequenced in parallel with total mRNA (Sup.2A). First, we compared polysome profiles of IL-1β treated and untreated RAC (Fig.1D, Sup.2B). Fold changes (FC) of mRNApoly ≥1.5 were significantly different for 369 genes (Sup.Table1). These changes can be mediated by changes in mRNA abundance, or/and by translational regulation. We therefore evaluated transcriptional changes in IL-1β treated RAC (Sup. 2C). Out of 12721 genes, 776 genes were differentially expressed (FC≥1.5) (Sup.Table2). RNAseq data were validated further with qPCR (R=0.84; Sup. 2D). For majority of identified genes mRNAs abundance determined their expression. Thus, IL-1β induced increase in total Mmp13 mRNA and mRNA_poly_ levels is similar (7.95 vs. 7.48), meaning that Mmp13 expression is regulated predominantly at the transcriptional level. Likewise, transcription determines monoglyceride Lipase (Mgll) expression, as both Mgll mRNA total and mRNA_poly_ were decreased (Fig.2A).

**Figure 2.**
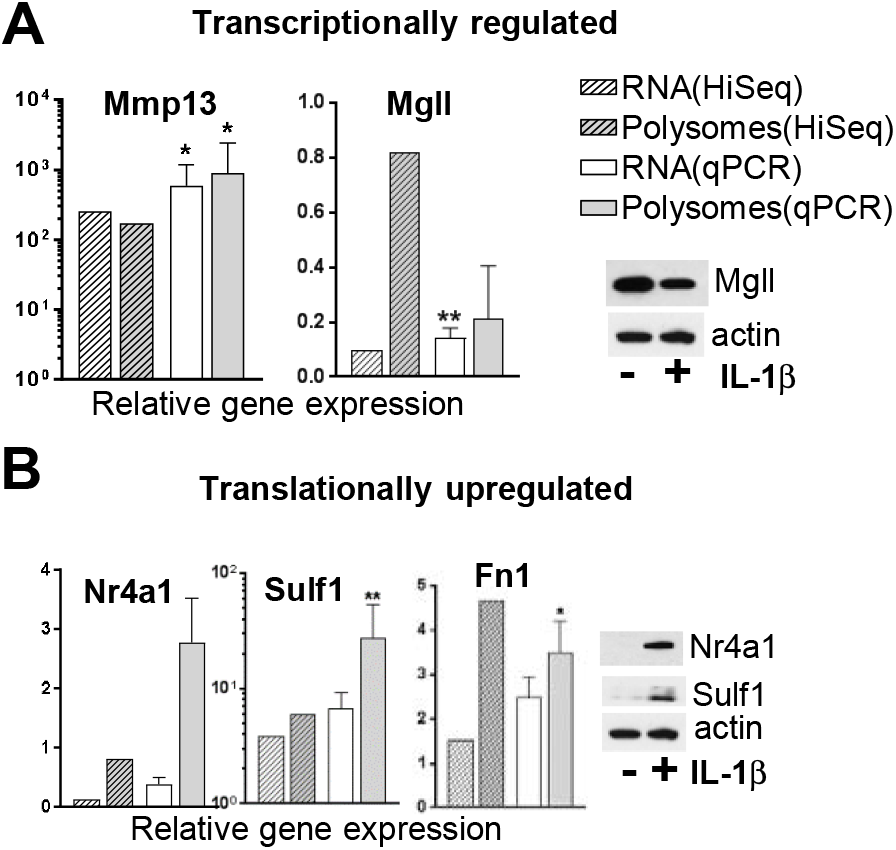
Validation of RNAseq data. Validation of **(A)** transcriptionally and **(B)** translationally regulated targets. Bars show levels of gene expression as a FC (treated vs untreated) (n=3, *t*-test,*p ≤ 0.05, **p ≤ 0.01). WB of IL1β-treated and untreated RAC confirms expression of the genes of interest; (α-actin is used as a loading control, 20-40μg of total protein, or 1/20 of a collected supernatant normalized to total protein).

To identify translationally regulated genes, we merged two datasets (mRNA_poly_ (369 genes) and total mRNA (776)) generating a set of 848 genes. 617 genes with the ratio between FC(Total mRNA(IL-1β/control)) to FC(mRNA_poly_(IL-1β/control) ≥1.3 and ≤0.77 (an arbitrary cutoff implying that there is a substantial translational component in the regulation of gene expression) were taken for further consideration. For example, for Fn1 and Sulfatase 1 (Sulf1), FCs between transcriptome and translatome were significantly different (0.54 vs 2.33) and (2.71 vs. 1.88) respectively (Fig.2B). 100 out of 617 entries that had RNAseq data for both total mRNA and mRNA_poly_ (in bold in Sup.Table4) were analyzed further. Top translationally upregulated by IL-1β signaling mRNAs include Anthrax Toxin Receptor2, Fn1, and Cathelicidin Antimicrobial Peptide (Sup.Table 5). Other examples of translationally upregulated genes include Plasminogen Activator, Urokinase (Plau) and Nuclear Receptor Subfamily 4 Group A Member 1 (Nr4a1) (Fig.2B, Sup.3A). Increased protein expression of Sulf1 (32, 33), Fn1 (2), Plau (34), but not Nr4a1 was previously reported in OA cartilage.

Importantly, Gene ontology analysis shows that translationally regulated genes are enriched for ECM genes (Sup.Table 6). This suggests the distinct role of translation control for ECM homeostasis in articular chondrocytes.

Remarkably, the polysome profiles of untreated RAC revealed that 1846 mRNAs (15.6% of all detected genes) are translationally repressed (LogFC(Total mRNA)/(mRNA_poly_)) ≥ 2.0) (Sup.Table7; Sup.2F). For instance, only minor portions of mRNAs encoding known chondrocytes markers Col2a1 and Acan were found on polysomes. The top 10 translationally repressed mRNAs include those encoding Fn1 (about 1% is being translated), thrombospondin and proteins involved in cytoskeleton formation. Among efficiently translated mRNAs in unchallenged chondrocytes we detected cartilage-specific protein CD-RAP/MIA, Phospholipase A2 Group IIA, and S100 Calcium Binding Protein A6 (35).

Thus, polysome profiling identified a substantial set of genes whose protein abundance is translationally regulated by IL-1β signaling.

### Expression of translationally activated targets in rodent and human OA cartilage

We next tested if identified targets are translationally regulated *in vivo*. Immunostaining (IHC) of Sulf1, Plau and Nr4a1 was significantly increased in operated compared to sham joints. The similar result was obtained when cells isolated from animals were analyzed by WB or IF (Fig.3A, 3D, Sup.3B). Importantly, no significant changes were observed in Nr4a1 and Fn mRNA levels isolated from either sham or ACLT joints (Sup.1E). We next analyzed lesional and non-lesional areas of human OA cartilage isolated from the same patient (n=4). We observed increased immunostaining in lesion areas for Sulf1, Nr4a1, Mmp13, and Plau (Fig.3B; Sup.3C) (36). No significant changes were detected in Nr4a1 mRNA level when OA (n=12) and non-OA (n=9) cartilage was analyzed by qPCR or by Affymetrix U133A microarray (Fig.3C; Sup.3D, 3E). Sulf1 mRNA expression were slightly upregulated in OA compared to normal cartilage (similar to our polysome profiling data) indicating that synergy between transcriptional and translational regulation drives its expression. We next treated cells isolated from non-lesion areas with IL-1β to confirm that translational upregulation is mediated by IL-1β signaling. Nr4a1 and Fn1 protein levels were increased (Fig.3E). The level of upregulation fluctuated from a patient to a patient, reflecting the heterogeneous nature of the diseases. Importantly, there were no significant changes in Fn1 and Nr4a1 mRNA levels (Sup.3F). Our data indicate that increased protein abundance of Fn1 and Nr4a1 is primarily controlled at the translational level in OA cartilage and likely mediated by pro-inflammatory signaling.

**Figure 3.**
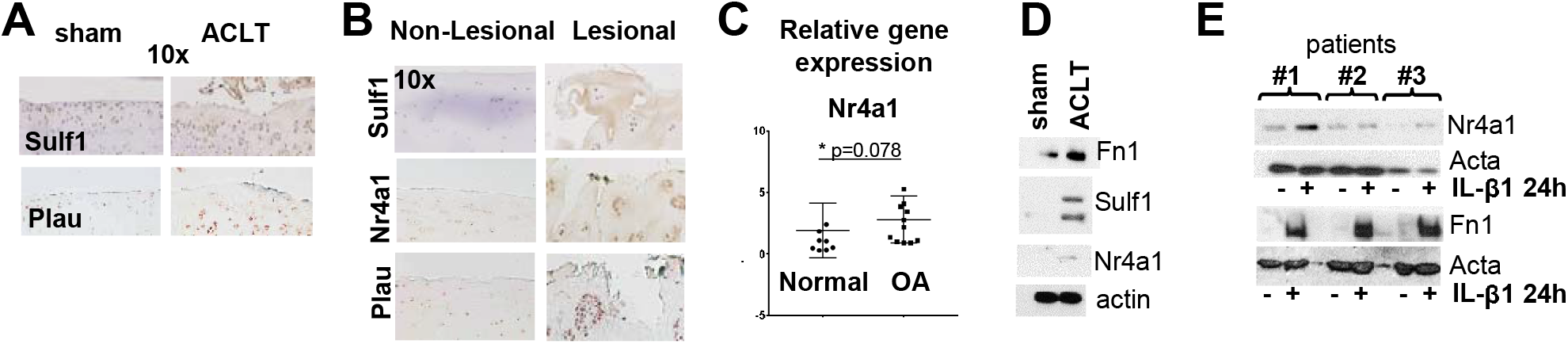
Expression of translationally controlled targets *in vivo*. **(A)** IHC analyses of indicated targets in the ACLT rat model at 20 weeks post-op. **(B)** Representative IHC analysis of non-lesional and lesional areas of cartilage from the same OA patient (n=5). **(C)** The graph shows levels of Nr4a1 mRNA expression, analyzed by qPCR as a FC (normal (n=9) vs OA (n=12) cartilage), Mann-Whitney test. **(D)** WB of cultured chondrocytes from ACLT and sham-operated knee; (α-actin is used as a loading control, loading 40μg of total protein. **(E)** WB of IL-1β-treated chondrocytes isolated from the non-lesional area of human cartilage (n=5; three representative samples are shown;α-actin is used as a loading control, 40μg of total protein).

#### 5’ Untranslated Regions (UTRs) are responsible for IL-1β mediated translational upregulation

We demonstrated that 4E-BP, an inhibitor of cap-dependent translation is inactivated in OA cartilage (18). 4E-BP acts as a repressor by sequestering eukaryotic initiation factor 4E (eIF4E) cap-binding protein, which is a part of eIF4F complexes. eIF4F is crucial for binding of the ribosomal complexes onto the 5’cap structure of mRNAs and unwinding RNA structures that precede to the initiation codon. It is well established that different mRNAs have different requirements for eIF4F depending on the length and structure of their 5’UTRs.

mTORC1-mediated phosphorylation of Thr37/46/70 and Ser65 restricts 4E-BP ability to bind eIF4E. We next determined if 4E-BP inactivation (which would ultimately increase the amount of active eIF4F) is crucial for upregulation of genes of our interest at the protein level. We used 4E-BP(AA) dominant-negative mutant which has Thr37/46 to Ala substitutions and therefore cannot be inactivated by phosphorylation (37). 4E-BP(AA) overexpression strongly reduced mRNA_poly_ and protein upregulation of examined targets when compared to RAC expressing a parental vector supporting our hypothesis that their 5’UTRs might play an important role in regulation of translation (Fig.4A;Sup.4A,4B). We next examined if Fn1, Sulf1, and Nr4a1 5’UTRs are functionally important using a luciferase reporter.

**Figure4.**
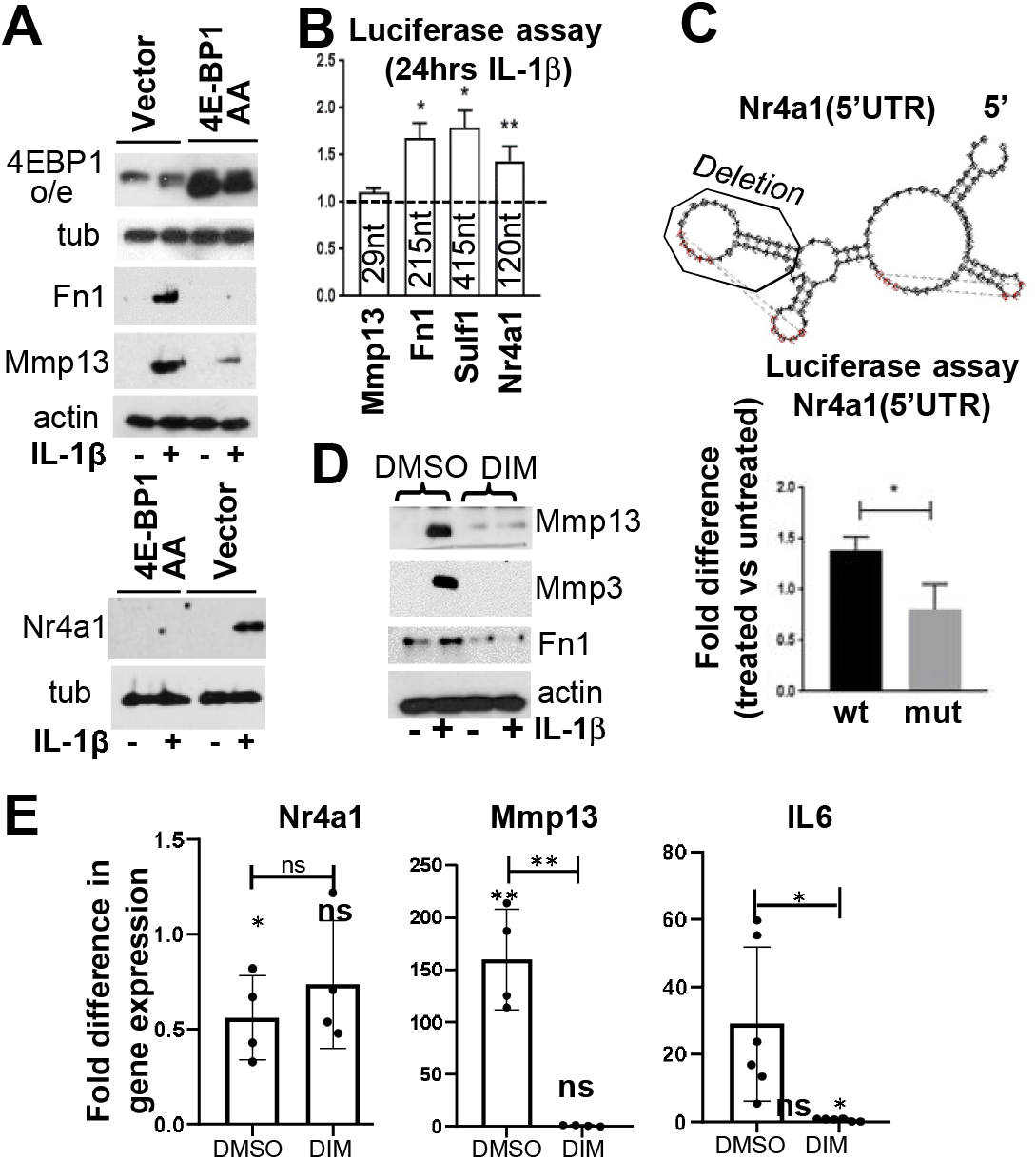
5’UTRs are responsible for IL-1β mediated translational upregulation. **(A)** Dominant-negative 4E-BP1(AA) mutant (or parental vector) was expressed in RAC. WB of IL-1β-treated RAC (α-actin and tubulin are used as a loading control, 40μg of total protein, or 1/20 of a collected supernatant normalized to total protein (Mmp13)). **(B)** Bars demonstrate 5’UTR activity in a luciferase reporter assay. Indicated 5’ UTRs were cloned into pIS0 plasmid. RAC were treated with IL-1β for 24hrs (n=3; data are represented as means±s.e.m. of fold difference in luminescence of treated vs. untreated cells; *t*-test, *p≤ 0.05, **p ≤ 0.01). **(C)** Mutational analysis of Nr4a1 5’UTR. The deletion is indicated on the predicted structure. A luciferase reporter assay was performed using wild-type (wt) and mutated (mut) 5’UTRs (n=3; *t*-test,*p ≤ 0.05). **(D,E) IL-1β mediated increase in Mmps expression requires Nr4a1**. Cells were pre-treated with DMSO or DIM, treated for 24hrs with IL1β and analyzed by **(D)** WB or **(E)** qPCR (n=4; data are represented as means±sd; significance is estimated for each condition individually as compared to untreated cells (indicated above each bar; unpaired t-test); significance between different conditions is indicated above the brackets (two-tailed paired t-test; *p≤ 0.05, **p ≤ 0.01).

5’UTRs were cloned into pIS0 vector upstream a luciferase cDNA, and their activity upon IL-1β stimulation was assayed. Mmp13 5’UTR was used as a negative control. An increase in Luc expression was detected for all but Mmp13 5’UTR (Fig.4B). While Fn1 and Sulf1 effect was anticipated, as their 5’UTRs are quite long (215 and 415nts), the strong effect of Nr4a1 5’UTR (120nt) was rather surprising. This sequence can potentially form a highly structured fold that includes pseudoknots (Fig.4C), thus increasing the demand in eIF4F. We disrupted this structure by deleting 27nts in the loop structure. The Nr4a1 5’UTR mutant failed to upregulate Luc expression upon IL-1β stimulation, indicating the importance of this element/structure for efficient Nr4a1 translation (Fig.4C). This result supports our hypothesis that 5’UTRs of Fn1, Sulf1, and Nr4a1 are important determinants of their effective translation and are sensitive to 4E-BP activity upon IL-1β signaling *in vitro* and likely *in vivo*. Other pro-inflammatory cytokines stimulating mTORC1 activity would ultimately lead to 4E-BP inactivation as well and have the similar effect on translationally regulated genes.

### IL-1β mediated increase in ECM metalloproteinases requires Nr4a1

To corroborate the importance of newly identified Nr4a1 for OA pathology, we investigated if its functions are important for mediating pro-inflammatory response in chondrocytes. Nr4a1 is an orphan nuclear transcription factor that is involved in multiple processes in metabolism, and inflammation (38). It was reported that Nr4a1 mRNA level is similar in OA and normal cartilage, and another member of this receptor family-Nr4a2 was implicated in synoviocyte invasion and Mmp13 expression (39). We inhibited Nr4a1 functions by the *p*-hydroxyphenyl analog [1,1-bis(3’-indolyl)-1-(*p*-hydroxyphenyl)methane (DIM-C-pPhOH)] (DIM here and in Fig.4) (40). Nr4a1 inhibition significantly prevented IL-1β induced Mmp13 and Mmp3 expression at mRNA and protein levels (Fig.4D,E;Sup.5). This robust response is likely due to suppression of IL-6 expression in IL-1β treated cells in the presence of DIM (Fig.4E), as Nr4a1 is IL-6 upstream regulator (41). IL-6, in turn, is linked to OA progression (42). Thus, Nr4a1 is important for mediating IL-6 expression and a pro-inflammatory response of chondrocytes. More experiments are needed in order to understand the mechanism of Nr4a1 functions in cartilage homeostasis starting with the characterization of the chondrocyte-specific repertoire of Nr4a1 targets.

### Inhibition of cap-binding transaltion delays OA progression in the ACLT model

mTORC1 is a master regulator of 4E-BP activity. mTORC1 inhibition attenuates the effect of IL-1β signaling on protein synthesis and MMPs expression in articular chondrocytes (18). Intraperitoneal administration of Rapamycin, a mTORC1 inhibitor (43), significantly reduced the severity of cartilage degradation in a mouse model of post-traumatic OA (44).The similar protective effect was observed with the cartilage-specific ablation of mTOR (45). In both cases, autophagy was assessed as a downstream target of mTOR, though 4E-BP should be affected as well. We specifically inhibited mTORC1/4E-BP/eIF4E axis in order to examine its role in OA progression. We used 4E1RCat inhibitor that blocks cap-dependent translation by inhibiting eIF4E (46). *In vitro*, 4E1RCat prevented IL-1β induced upregulation of protein synthesis and Mmps in RAC (Fig.5A,5B,Sup.6A,6B). 4E1RCat (or a vehicle) was injected into ACLT or sham-operated joints (n=7 for each group) 8 weeks post-op weekly for 4 weeks (Sup. 6C). This timing was based on our data that upregulation of total protein synthesis is detected at 12 weeks post-op in our OA model 4E1RCat treatment significantly reduced cartilage degeneration in operating joints (Fig.5C). Consistent with these data, the level of active Mmp13 was also significantly reduced in cells isolated from ACLT joints treated with 4E1RCat (Fig.5D). Sulf1, Nr4a1, and Fn1 protein abundance was decreased in the 4E1RCat treated joints as analyzed by WB and IHC (Fig.5E,Sup.6D). Thus, intra-articular administration of the eIF4E inhibitor reduced cartilage degeneration in OA joints and attenuated the effect of injury on the expression of translationally controlled genes.

**Figure5.**
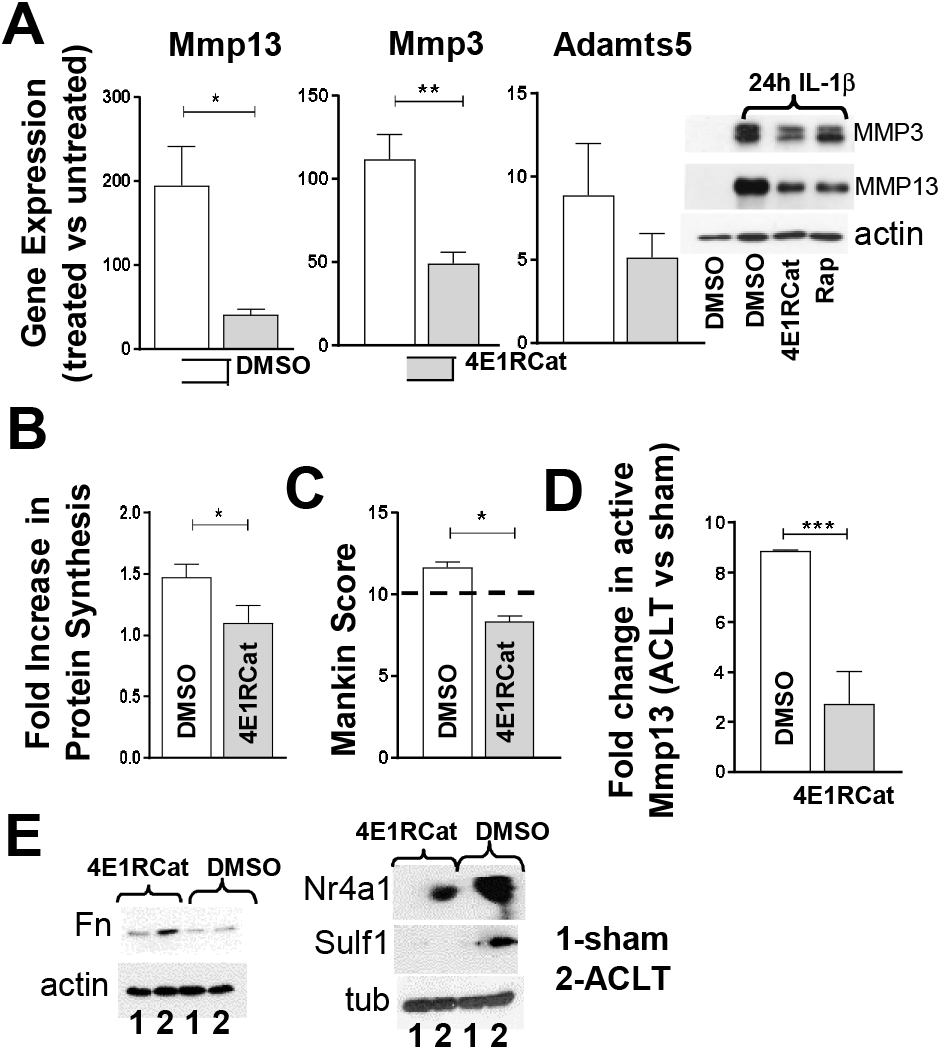
Inhibition of cap-binding translation delays OA progression *in vivo*. **(A)** RAC were pretreated with 4E1Rcat (10uM) for 1hr followed by 24hrs of IL1β (10nM). mRNA expression was analyzed by qPCR (n=3; data are represented as treated vs. untreated; student’s t-test, *p≤ 0.05, **p ≤ 0.01). WB confirms Mmp13 and Mmp3 downregulation (15ug of total protein, actin used as a loading control, or 1/20 of a collected supernatant normalized to total protein). **(B)** The total rate of protein synthesis was significantly decreased in the presence of 4E1Rcat in IL1β treated RAC. Bars demonstrate FC in protein synthesis with or without 4E1Rcat (n=3; data are represented as means±s.e.m.; student’s t-test, *p ≤ 0.05). Disease progression is monitored 12 weeks post-op using Modified Mankin score **(C)** and changes in active Mmp13 levels **(D)** (n=4; scoring by two independent users, student’s t-test, *p≤ 0.05). The line shows severe disease level. Bars represents FC of active Mmp13 levels in ACLT vs. sham in the vehicle compared to 4E1Rcat treatment (n=3, student’s t-test, ***p ≤ 0.001). **(E)** WB analyses of cultured chondrocytes from ACLT and sham-operated knee treated with 4E1RCat or vehicle; (n=3; α-actin and tubulin are used as a loading control, 40μg of total protein).

## DISCUSSION

The presented results provide evidence for an important role of translational control in cartilage homeostasis. As our polysome profiling data are unique, we can only compare them with existing transcriptome data of pro-inflammatory response in cartilage. The observed similarities include increased expression of numerous inflammatory mediators (IL-6, IL23r, and CXCL1), and growth factors (BMP-2 and BMP-6) (47, 48). Aggrecan-1 downregulation and increase in several MMPs expression were also detected.

Polysome profiling revealed that many components of ECM including fibronectin are translationally repressed in unchallenged chondrocytes despite relatively high mRNA levels. This repression is consistent with the low mitotic activity of the residing chondrocytes (49, 50). However, high mRNA abundance of Fn1 in unchallenged chondrocytes makes this gene a quick responder to different stimulus through protein abundance in OA cartilage. Interestingly, while human FN1 gene has three alternatively spliced regions, with the potential to produce 20 different transcript variants, most of these variants have the same 5’UTR which as we demonstrated is important for Fn1 efficient translation. Our data are in line with an early work that suggested the regulation of Fn1 expression in OA cartilage through protein synthesis (2) as well as with data demonstrated an increase in Fn1 protein (but not mRNA) level in OA cartilage (10, 11, 13). It is important to notice that while we used IL-1β, other pro-inflammatory cytokines stimulating mTORC1 activity would ultimately lead to 4E-BP inactivation and have similar effect on translationally regulated genes and on protein synthesis in general. Further work is needed to understand the specific contribution of diverse signaling pathways involved in OA pathology (51). We demonstrated that in post-traumatic animal OA model 4E-BP inactivation in articular cartilage is one of the earliest events following traumatic insult onto the joint that precedes Mmp13 and Mmp3 upregulation and degeneration of articular cartilage (52). Immediate 4E-BP inactivation in the post-traumatic joint is likely advantageous for a specific pool of mRNAs those translational efficiencies strongly depend on eIF4F activity, such as identified here Fn1 and Nr4a1. The dependence of translational landscape on 4E-BP/eIF4F activity likely changes as disease advances, adjusting to epigenetic and metabolic changes resulted from mechanical stress and low grade inflammation processes.

Polysome analysis of IL-1β treated articular chondrocytes revealed novel potential players in OA pathology, such as an orphan nuclear receptor Nr4a1 those abundance is elevated in human and rodent OA cartilage at protein but not mRNA level. Mix et al. also reported that Nr4a1 mRNA levels are similar in OA and normal cartilage (53). Our experiments with DIM inhibitor demonstrated that Nr4a1 is crucial for mediating IL-6 expression and a pro-inflammatory response of chondrocytes in general. More experiments are needed in order to understand the mechanism of Nr4a1 functions in cartilage homeostasis starting with the characterization of the chondrocyte-specific repertoire of Nr4a1 targets. Our *in vitro* and *in vivo* data provided a rationale for targeting translational apparatus to limit/delay the progression of cartilage degeneration in OA. In support of our premises, genetic or pharmacological inhibition of mTORC1 delayed cartilage degeneration in a mouse model of post-traumatic OA (44, 45). To target protein synthesis specifically, we used 4E1RCat, a chemical inhibitor of eIF4E. Intra-articular administration of 4E1RCat reduced cartilage degeneration and attenuated the effect of ACLT on the expression of translationally controlled genes: Fn1, Sulf1, and Nr4a1, but not actin. With further optimization of regime and delivery approach, 4E1RCat (or similar compounds) might be considered for clinical applications.

Another important aspect of post-transcriptional control - RNA/protein turnover should also be assessed in future studies. Thus, it was demonstrated that the stability of Plau mRNA is increased in OA tissue, making mRNA turnover an additional important factor that regulates Plau expression (16). We demonstrated that Plau protein abundance is also translationally upregulated upon pro-inflammatory signaling. Thus, this example shows how multiple levels of regulation converge to adapt to tissue specific changes.

In summary, using polysome profiling, we identified novel OA-related targets, thus underlying the exceptional potential of our approach for discovering new players in human OA cartilage. These functionally unrelated targets might be regulated as a selective pool by fine-tuning their translation efficiency through careful incremental modulation of eIF4F activity, while keeping translation of most of the genes unchanged.

## ACKNOWLEDGMENTS

We are grateful to Dr. Andreev and Dr. Mignatti for the critical reviewing of the manuscript. This work was supported by NIH grants AR063128 (to V.K.) and in part R01-AR054817 (to S.B.A.).

## CONTRIBUTORSHIP

Conceptualization, V.K; Methodology, O.K., V.K., and M.A; Investigation, O.K., V.K., and M.A; Writing – Original Draft, O.K., V.K., and M.A; Visualization, O.K., V.K., and M.A Funding Acquisition, and V.K.; Resources, V.K., and M.A; Supervision, V.K. and M.A.

## ETHICAL APPROVAL INFORMATION

All animal experiments were performed in accordance with the regulations of the Institutional Animal Care and Use Committee at NYU School of Medicine.

## DATA SHARING

All data are included in the main text or as supplemental material.

## REFERENCES

1. Brown RA, Jones KL. The synthesis and accumulation of fibronectin by human articular cartilage. J Rheumatol. 1990;17(1):65–72.

2. Lust G, Burton-Wurster N, Leipold H. Fibronectin as a marker for osteoarthritis. J Rheumatol. 1987;14 Spec No:28–9.

3. Stoffels JM, Zhao C, Baron W. Fibronectin in tissue regeneration: timely disassembly of the scaffold is necessary to complete the build. Cell Mol Life Sci. 2013;70(22):4243–53.

4. Ding L, Guo D, Homandberg GA. Fibronectin fragments mediate matrix metalloproteinase upregulation and cartilage damage through proline rich tyrosine kinase 2, c-src, NF-kappaB and protein kinase Cdelta. Osteoarthritis Cartilage. 2009;17(10):1385–92.

5. Zack MD, Arner EC, Anglin CP, Alston JT, Malfait AM, Tortorella MD. Identification of fibronectin neoepitopes present in human osteoarthritic cartilage. Arthritis Rheum. 2006;54(9):2912–22.

6. Forsyth CB, Pulai J, Loeser RF. Fibronectin fragments and blocking antibodies to alpha2beta1 and alpha5beta1 integrins stimulate mitogen-activated protein kinase signaling and increase collagenase 3 (matrix metalloproteinase 13) production by human articular chondrocytes. Arthritis Rheum. 2002;46(9):2368–76.

7. Mimura Y, Ihn H, Jinnin M, Asano Y, Yamane K, Tamaki K. Epidermal growth factor induces fibronectin expression in human dermal fibroblasts via protein kinase C delta signaling pathway. J Invest Dermatol. 2004;122(6):1390–8.

8. Ramos YF, den Hollander W, Bovee JV, Bomer N, van der Breggen R, Lakenberg N, et al. Genes involved in the osteoarthritis process identified through genome wide expression analysis in articular cartilage; the RAAK study. PLoS One. 2014;9(7):e103056.

9. Fu M, Huang G, Zhang Z, Liu J, Zhang Z, Huang Z, et al. Expression profile of long noncoding RNAs in cartilage from knee osteoarthritis patients. Osteoarthritis Cartilage. 2015;23(3):423–32.

10. Loeser RF, Olex AL, McNulty MA, Carlson CS, Callahan MF, Ferguson CM, et al. Microarray analysis reveals age-related differences in gene expression during the development of osteoarthritis in mice. Arthritis Rheum. 2012;64(3):705–17.

11. Zhu W, Wang D, Lu W, Han Y, Ou Y, Zhou K, et al. Gene expression profile of the synovium and cartilage in a chronic arthritis rat model. Artif Cells Blood Substit Immobil Biotechnol. 2012;40(1-2):70–4.

12. Wei T, Kulkarni NH, Zeng QQ, Helvering LM, Lin X, Lawrence F, et al. Analysis of early changes in the articular cartilage transcriptisome in the rat meniscal tear model of osteoarthritis: pathway comparisons with the rat anterior cruciate transection model and with human osteoarthritic cartilage. Osteoarthritis Cartilage. 2010;18(7):992–1000.

13. Kuttapitiya A, Assi L, Laing K, Hing C, Mitchell P, Whitley G, et al. Microarray analysis of bone marrow lesions in osteoarthritis demonstrates upregulation of genes implicated in osteochondral turnover, neurogenesis and inflammation. Ann Rheum Dis. 2017;76(10):1764–73.

14. Snelling S, Rout R, Davidson R, Clark I, Carr A, Hulley PA, et al. A gene expression study of normal and damaged cartilage in anteromedial gonarthrosis, a phenotype of osteoarthritis. Osteoarthritis Cartilage. 2014;22(2):334–43.

15. Vogel C, Marcotte EM. Insights into the regulation of protein abundance from proteomic and transcriptomic analyses. Nat Rev Genet. 2012;13(4):227–32.

16. Tew SR, McDermott BT, Fentem RB, Peffers MJ, Clegg PD. Transcriptome-wide analysis of messenger RNA decay in normal and osteoarthritic human articular chondrocytes. Arthritis Rheumatol. 2014;66(11):3052–61.

17. McDermott BT, Ellis S, Bou-Gharios G, Clegg PD, Tew SR. RNA binding proteins regulate anabolic and catabolic gene expression in chondrocytes. Osteoarthritis Cartilage. 2016;24(7):1263–73.

18. Katsara O, Attur M, Ruoff R, Abramson SB, Kolupaeva V. Increased Activity of the Chondrocyte Translational Apparatus Accompanies Osteoarthritic Changes in Human and Rodent Knee Cartilage. Arthritis Rheumatol. 2017;69(3):586–97.

19. Le LT, Swingler TE, Clark IM. Review: the role of microRNAs in osteoarthritis and chondrogenesis. Arthritis Rheum. 2013;65(8):1963–74.

20. Tetlow LC, Adlam DJ, Woolley DE. Matrix metalloproteinase and proinflammatory cytokine production by chondrocytes of human osteoarthritic cartilage: associations with degenerative changes. Arthritis Rheum. 2001;44(3):585–94.

21. Favero M, Belluzzi E, Trisolino G, Goldring MB, Goldring SR, Cigolotti A, et al. Inflammatory molecules produced by meniscus and synovium in early and end-stage osteoarthritis: a coculture study. J Cell Physiol. 2018.

22. Watson Levings RS, Broome TA, Smith AD, Rice BL, Gibbs EP, Myara DA, et al. Gene Therapy for Osteoarthritis: Pharmacokinetics of Intra-Articular Self-Complementary Adeno-Associated Virus Interleukin-1 Receptor Antagonist Delivery in an Equine Model. Hum Gene Ther Clin Dev. 2018;29(2):90–100.

23. Elsaid KA, Ubhe A, Shaman Z, D’Souza G. Intra-articular interleukin-1 receptor antagonist (IL1-ra) microspheres for posttraumatic osteoarthritis: in vitro biological activity and in vivo disease modifying effect. J Exp Orthop. 2016;3(1):18.

24. Stone A, Grol MW, Ruan MZC, Dawson B, Chen Y, Jiang MM, et al. Combinatorial Prg4 and Il-1ra Gene Therapy Protects Against Hyperalgesia and Cartilage Degeneration in Post-Traumatic Osteoarthritis. Hum Gene Ther. 2018.

25. Watson Levings RS, Smith AD, Broome TA, Rice BL, Gibbs EP, Myara DA, et al. Self-Complementary Adeno-Associated Virus-Mediated Interleukin-1 Receptor Antagonist Gene Delivery for the Treatment of Osteoarthritis: Test of Efficacy in an Equine Model. Hum Gene Ther Clin Dev. 2018;29(2):101–12.

26. Nixon AJ, Grol MW, Lang HM, Ruan MZC, Stone A, Begum L, et al. Disease-Modifying Osteoarthritis Treatment With Interleukin-1 Receptor Antagonist Gene Therapy in Small and Large Animal Models. Arthritis Rheumatol. 2018;70(11):1757–68.

27. Piccirillo CA, Bjur E, Topisirovic I, Sonenberg N, Larsson O. Translational control of immune responses: from transcripts to translatomes. Nat Immunol. 2014;15(6):503–11.

28. Attur MG, Palmer GD, Al-Mussawir HE, Dave M, Teixeira CC, Rifkin DB, et al. F-spondin, a neuroregulatory protein, is up-regulated in osteoarthritis and regulates cartilage metabolism via TGF-beta activation. FASEB J. 2009;23(1):79–89.

29. Homandberg GA, Hui F, Wen C, Purple C, Bewsey K, Koepp H, et al. Fibronectin-fragment-induced cartilage chondrolysis is associated with release of catabolic cytokines. Biochem J. 1997;321 (Pt 3):751–7.

30. Gerwin N, Bendele AM, Glasson S, Carlson CS. The OARSI histopathology initiative - recommendations for histological assessments of osteoarthritis in the rat. Osteoarthritis Cartilage. 2010;18 Suppl 3:S24–34.

31. Long DL, Loeser RF. p38gamma mitogen-activated protein kinase suppresses chondrocyte production of MMP-13 in response to catabolic stimulation. Osteoarthritis Cartilage. 2010;18(9):1203–10.

32. Otsuki S, Taniguchi N, Grogan SP, D’Lima D, Kinoshita M, Lotz M. Expression of novel extracellular sulfatases Sulf-1 and Sulf-2 in normal and osteoarthritic articular cartilage. Arthritis Res Ther. 2008;10(3):R61.

33. Goldring MB. Chondrogenesis, chondrocyte differentiation, and articular cartilage metabolism in health and osteoarthritis. Ther Adv Musculoskelet Dis. 2012;4(4):269–85.

34. Tang YL, Zhu GQ, Hu L, Zheng M, Zhang JY, Shi ZD, et al. Effects of intra-articular administration of sodium hyaluronate on plasminogen activator system in temporomandibular joints with osteoarthritis. Oral Surg Oral Med Oral Pathol Oral Radiol Endod. 2010;109(4):541–7.

35. Bosserhoff AK, Kondo S, Moser M, Dietz UH, Copeland NG, Gilbert DJ, et al. Mouse CD-RAP/MIA gene: structure, chromosomal localization, and expression in cartilage and chondrosarcoma. Dev Dyn. 1997;208(4):516–25.

36. Geyer M, Grassel S, Straub RH, Schett G, Dinser R, Grifka J, et al. Differential transcriptome analysis of intraarticular lesional vs intact cartilage reveals new candidate genes in osteoarthritis pathophysiology. Osteoarthritis Cartilage. 2009;17(3):328–35.

37. Gingras AC, Raught B, Gygi SP, Niedzwiecka A, Miron M, Burley SK, et al. Hierarchical phosphorylation of the translation inhibitor 4E-BP1. Genes Dev. 2001;15(21):2852–64.

38. Lee SO, Li X, Khan S, Safe S. Targeting NR4A1 (TR3) in cancer cells and tumors. Expert Opin Ther Targets. 2011;15(2):195–206.

39. Mix KS, McMahon K, McMorrow JP, Walkenhorst DE, Smyth AM, Petrella BL, et al. Orphan nuclear receptor NR4A2 induces synoviocyte proliferation, invasion, and matrix metalloproteinase 13 transcription. Arthritis Rheum. 2012;64(7):2126–36.

40. Lee SO, Abdelrahim M, Yoon K, Chintharlapalli S, Papineni S, Kim K, et al. Inactivation of the orphan nuclear receptor TR3/Nur77 inhibits pancreatic cancer cell and tumor growth. Cancer Res. 2010;70(17):6824–36.

41. Wang SC, Myers SA, Eriksson NA, Fitzsimmons RL, Muscat GE. Nr4a1 siRNA expression attenuates alpha-MSH regulated gene expression in 3T3-L1 adipocytes. Mol Endocrinol. 2011;25(2):291–306.

42. Appleton CT. Osteoarthritis year in review 2017: biology. Osteoarthritis Cartilage. 2018;26(3):296–303.

43. Ballou LM, Lin RZ. Rapamycin and mTOR kinase inhibitors. J Chem Biol. 2008;1(1-4):27–36.

44. Carames B, Hasegawa A, Taniguchi N, Miyaki S, Blanco FJ, Lotz M. Autophagy activation by rapamycin reduces severity of experimental osteoarthritis. Ann Rheum Dis. 2012;71(4):575–81.

45. Zhang Y, Vasheghani F, Li YH, Blati M, Simeone K, Fahmi H, et al. Cartilage-specific deletion of mTOR upregulates autophagy and protects mice from osteoarthritis. Ann Rheum Dis. 2015;74(7):1432–40.

46. Moerke NJ, Aktas H, Chen H, Cantel S, Reibarkh MY, Fahmy A, et al. Small-molecule inhibition of the interaction between the translation initiation factors e1F4E and e1F4G. Cell. 2007;128(2):257–67.

47. Gouze JN, Bianchi A, Becuwe P, Dauca M, Netter P, Magdalou J, et al. Glucosamine modulates IL-1-induced activation of rat chondrocytes at a receptor level, and by inhibiting the NF-kappa B pathway. FEBS Lett. 2002;510(3):166–70.

48. Fukui N, Zhu Y, Maloney WJ, Clohisy J, Sandell LJ. Stimulation of BMP-2 expression by pro-inflammatory cytokines IL-1 and TNF-alpha in normal and osteoarthritic chondrocytes. J Bone Joint Surg Am. 2003;85-A Suppl 3:59–66.

49. Gomez-Camarillo MA, Kouri JB. Ontogeny of rat chondrocyte proliferation: studies in embryo, adult and osteoarthritic (OA) cartilage. Cell Res. 2005;15(2):99–104.

50. Coburn JM, Bernstein N, Bhattacharya R, Aich U, Yarema KJ, Elisseeff JH. Differential response of chondrocytes and chondrogenic-induced mesenchymal stem cells to C1-OH tributanoylated N-acetylhexosamines. PLoS One. 2013;8(3):e58899.

51. Goldring SR, Goldring MB. The role of cytokines in cartilage matrix degeneration in osteoarthritis. Clin Orthop Relat Res. 2004(427 Suppl):S27–36.

52. Katsara O, Kolupaeva V. mTOR-mediated inactivation of 4E-BP1, an inhibitor of translation, precedes cartilage degeneration in rat osteoarthritic knees. J Orthop Res. 2018;36(10):2728–35.

53. Mix KS, Attur MG, Al-Mussawir H, Abramson SB, Brinckerhoff CE, Murphy EP. Transcriptional repression of matrix metalloproteinase gene expression by the orphan nuclear receptor NURR1 in cartilage. J Biol Chem. 2007;282(13):9492–504.

