## Supplemental File for "Control of cartilage homeostasis and osteoarthritis progression by mTORC1/4E-BP/eIF4E axis"

**SUPPLEMENTARY**

**METHODS**

*Cells culture conditions*. RAC were isolated and propagated as previously described ^1^. Briefly, cells were cultured in Chondrocyte Growth Medium (Cell applications) at 37°C and 5% CO_2_ until confluency, when experiments were performed. Cells were plated (100,000/cm^2^) and starved for 16 hours. For cap-dependent inhibition of translation cells were pretreated for 1hr with 4E1Rcat inhibitor (10uM or lower, ApexBio) prior to IL-1b (10 ng/ml) treatment*.*

*Isolation of translationally active pool of mRNAs by polysome fraction analysis.* RAC were expanded to 70% confluence. Then they were plated, starved for 16h and treated with 10 ng/ml of IL-1β for 24hrs. For the 10 final minutes cells were incubated with Cyclohexamide (100ug/ml) at 37^o^C. Cells were collected, washed twice with ice-cold PBS supplemented with Cyclohexamide (100ug/ml) and lysed on ice for 10min in buffer “A” (5mM TrisHCl, pH 7.5, 1.5mM KCl, 2.5mM MgCl_2_, protease EDTA-free cocktail (Sigma), 100ug/ml cyclohexamide, 2mM DTT, 0.5% Triton X-100, 0.5% Na deoxycholate, 50U RNase inhibitor). 2OD units of extract (3 independent experiments combined) were layered over 5 to 50% linear sucrose gradients in buffer “B” (20 mM HEPES, pH 7.5, 100mM KCl, 5 mM MgCl_2_, 100ug/ml cyclohexamide, protease EDTA-free cocktail (Sigma), RNase inhibitor) and centrifuged at 38,000 rpm for 2h at 4°C. Gradients were fractionated while monitoring RNA absorbance at 254nm. RNA from the heavy polysomes fractions was extracted with one volume of acid phenol–chloroform, and precipitated with one volume of isopropanol. RNA pellets were centrifuged at max speed for 20 min at 4°C, washed with 70% ethanol, and resuspended in 100ul of RNase-free water. Samples were further purified using Zymo Research RNA Clean and Concentrator-5, and eluted in 20ul RNase-free water. In parallel, cells were plated in 6 well plate (200,000 cells/well) for RNA and protein extraction. Sequencing of polysome bound RNA and total RNA was performed in parallel. Part of the sample was kept for validation by qPCR.

*Library preparation and sequencing*. Bioanalyzer (Agilent Technologies) and the RNA 6000 Nano Kit (Agilent Technologies) were used for size determination and quality control of the RNA. TruSeq Stranded Total RNA Library Preparation Kit with Ribo-Zero Gold (Illumina) was used to prepare the barcoded libraries from 100ng total RNA (12 amplification cycles). Libraries were validated and quantified using Bioanalyzer (Agilent); 11.5pM denatured libraries were used for deep sequencing (HiSeq 2500, Illumina) for 50 cycles.

*Downstream analysis.* An initial check of the raw reads was performed using FasQC v0.11.3 Reads containing adapter sequences and/or low quality ends were identified and trimmed using bbduc 35.59. A second quality check was performed to assure high-quality reads for the mapping step. Preprocedded reads were mapped to the Rat genome (Genome Assembly: Rnor6 (Ensembl release 89)) using NGS mapper STAR v2.5.2b ^2^. The alignment results of mapped reads are subsequently merged, and a sorted and indexed BAM file is created. Finally, mapping statistics files were generated. The alignments were overlapped with gene models using featureCounts v1.4.6-p5 ^3^. Multiple mappings and ambiguous hits were discarded to increase the reliability of the downstream analysis. Read counts were normalized by the absolute number of mapped reads (RPM) and by gene length (RPKM). Differentially expressed analysis was performed across all pair-wise combinations of conditions (resulting in 4 group comparison in total) using Bioconductor package egdeR v3.14.0 ^4^. Genes with a false discovery rate (FDR) less than 0.01 were considered to be significantly differentially expressed. Correlation analysis was performed using Rstudio. Enrichment analyses of biological processes were performed with DAVID (<https://david.ncifcrf.gov/>) (classification stringency – medium).

*qPCR analysis*. 500 ng of RNA was used for cDNA synthesis ( iScript cDNA synthesis kit, Bio-Rad). Quantitative PCR was run in a MyiQ Single-Color Real-Time PCR Detection System (Bio-Rad) using iQ SYBR Green Supermix. Relative gene expression levels were calculated using the ΔΔC_t_ method. Primers used are listed on Sup. Table 12.

*Transfection of RAC with Luc reporters*. To measure the translation of Luc reporter, RAC (10^5^ cells) were transfected with 2μg of each plasmid in 0.1 ml of Optimem containing 6μl Fugene6 (Promega, Madison,WI) for 24hrs. 24hrs letter cells were starved in DMEM/F12 containing 0.1% serum, and then treated with 10 ng/ml of IL-1β for 24hrs. Firefly Luc activity was measured using a Luciferase assay system (Promega, Madison,WI) according to the manufacturer’s protocol. Firefly luciferase activity was normalized to untreated control. Each assay was done in triplicate, and the mean and standard error were determined. Transfections with Nr4a1 siRNA (nt1701; Sigma) were performed using Lipofectamine RNAiMax according to the manufacturer's protocol. MISSION siRNA Universal Negative Control was used as a control. Cells were starved for 16h in serum-free medium 24h post-transfection and then treated with IL1-β.

*Experimental osteoarthritis in rats.* All animal experiments were performed according to IACUC regulations at NYU Langone Medical Center. Experimental osteoarthritis was induced in 12 week old male Sprague Dawley rats by transection of the anterior cruciate ligament in the right knee as described in ^5^. Knee joints from rats with experimental osteoarthritis (n=4) and sham-operated animals (n=4), were harvested. The joints were fixed in 4% PFA for 24h, decalcified in Formical2000 (Decal Chemical Corp, Tallman, NY, USA) for 96h, followed by paraffin embedding. Serial sections (4μm) were cut, stained with Safranin O-fast green and Toluidine Blue, and examined for histopathological changes using a semiquantitative scoring system.

*Intra-articular injections of 4E1Rcat inhibitor*. This method of drug delivery was chosen in order to minimize any possible adverse systemic side-effects of the drug (Larsen, C. et al. J. Pharm. Sci. 97, 4622–4654, 2008). Upon ACLT or sham operations, described elsewhere, all animals were allowed to move freely in plastic cages and had access to food and water ad libitum until they were sacrificed. To have a >90% power to detect a 70% reduction in OARSI score using an analysis of variance with repeated measures, with an α error probability of 0.05 and one group, we will need a minimum of four animals for each condition. Rats were treated with intra-articular injections of 100 μl of 4E1Rcat inhibitor (15 mg kg−1), or with DMSO in saline for 4 weeks starting week 8 post-op. Injections were performed once weekly in both groups. Rats were sacrificed 12 weeks post-op for IHC analysis.

*Immunohistochemical and IF analyses.* IHC and IF analysis of rat articular cartilage was performed as described previously ^1^. Briefly, the knee joints from ACLT-operated rats (n = 4) and sham-operated rats (n = 4) were harvested and fixed in 4% paraformaldehyde for 24 hours and then decalcified in Formical2000 (Decal Chemical) for 96 hours, followed by paraffin embedding. Serial sections (4μm) were stained with toluidine blue and Safranin O staining and examined for histopathologic changes, using a semiquantitative scoring system. For antigen retrieval, sections were incubated with proteinase K for 20 minutes at 37°C, followed by incubation with 0.2%Triton X-100 for 15 minutes. An avidin–biotin–peroxidase system (Vector) was used, and antigen–antibody complexes were visualized using diaminobenzidine as substrate. Tissue samples were counterstained with hematoxylin. Antibodies used are listed in Sup.Table 10. For IF analysis a secondary Cy5 fluorescence Ab was used. The nucleus was counterstained with DAPI.

*Human normal and OA cartilage microarray analysis***:** Was done as describedin ^6^. Total RNA from cartilage was isolated as previously described and 5 ug of total RNA was used for target preparation, and hybridized against the human U133A array according to the manufacturer’s instructions, and the expression levels were normalized as described previously ^6^.

1	Katsara, O., Attur, M., Ruoff, R., Abramson, S. B. & Kolupaeva, V. Increased Activity of the Chondrocyte Translational Apparatus Accompanies Osteoarthritic Changes in Human and Rodent Knee Cartilage. *Arthritis Rheumatol* **69**, 586-597, doi:10.1002/art.39947 (2017).

2	Dobin, A. *et al.* STAR: ultrafast universal RNA-seq aligner. *Bioinformatics* **29**, 15-21, doi:10.1093/bioinformatics/bts635 (2013).

3	Liao, Y., Smyth, G. K. & Shi, W. featureCounts: an efficient general purpose program for assigning sequence reads to genomic features. *Bioinformatics* **30**, 923-930, doi:10.1093/bioinformatics/btt656 (2014).

4	Robinson, M. D., McCarthy, D. J. & Smyth, G. K. edgeR: a Bioconductor package for differential expression analysis of digital gene expression data. *Bioinformatics* **26**, 139-140, doi:10.1093/bioinformatics/btp616 (2010).

5	Williams, J. M., Felten, D. L., Peterson, R. G. & O'Connor, B. L. Effects of surgically induced instability on rat knee articular cartilage. *J Anat* **134**, 103-109 (1982).

6	Attur, M. G. *et al.* F-spondin, a neuroregulatory protein, is up-regulated in osteoarthritis and regulates cartilage metabolism via TGF-beta activation. *FASEB J* **23**, 79-89, doi:10.1096/fj.08-114363 (2009).
