## Supplemental Figures for "Control of cartilage homeostasis and osteoarthritis progression by mTORC1/4E-BP/eIF4E axis"

### Slide 1
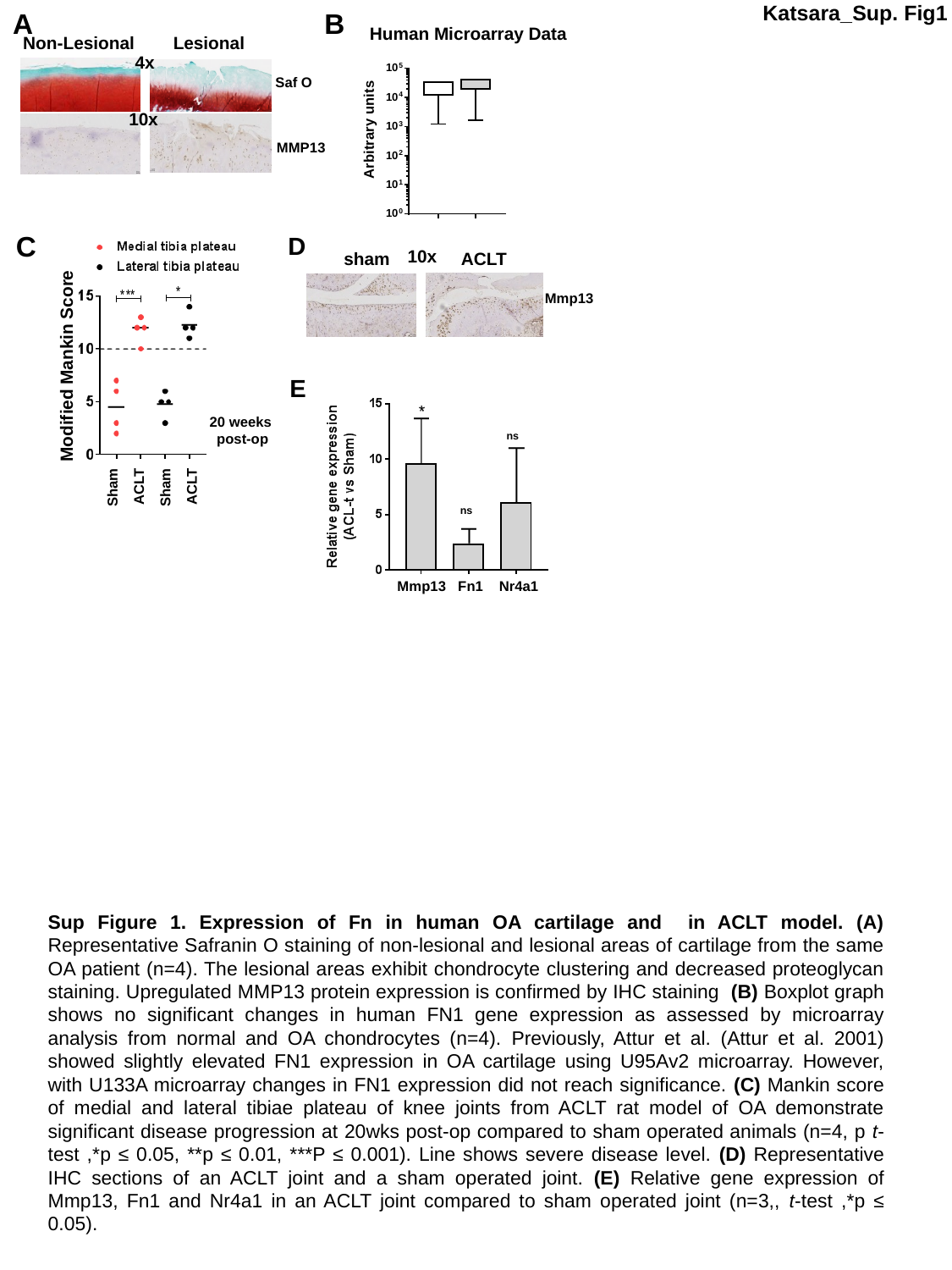

A
B
Katsara_Sup. Fig1
Non-Lesional Lesional
Human Microarray Data
4x
Saf O
Arbitrary units
10x
MMP13
C
D
Modified Mankin Score
20 weeks
post-op
Sham
ACLT
Sham
ACLT
10x
sham
ACLT
Mmp13
E
ns
ns
Mmp13 Fn1 Nr4a1
Sup Figure 1. Expression of Fn in human OA cartilage and in ACLT model. (A) Representative Safranin O staining of non-lesional and lesional areas of cartilage from the same OA patient (n=4). The lesional areas exhibit chondrocyte clustering and decreased proteoglycan staining. Upregulated MMP13 protein expression is confirmed by IHC staining (B) Boxplot graph shows no significant changes in human FN1 gene expression as assessed by microarray analysis from normal and OA chondrocytes (n=4). Previously, Attur et al. (Attur et al. 2001) showed slightly elevated FN1 expression in OA cartilage using U95Av2 microarray. However, with U133A microarray changes in FN1 expression did not reach significance. (C) Mankin score of medial and lateral tibiae plateau of knee joints from ACLT rat model of OA demonstrate significant disease progression at 20wks post-op compared to sham operated animals (n=4, p t-test ,*p ≤ 0.05, **p ≤ 0.01, ***P ≤ 0.001). Line shows severe disease level. (D) Representative IHC sections of an ACLT joint and a sham operated joint. (E) Relative gene expression of Mmp13, Fn1 and Nr4a1 in an ACLT joint compared to sham operated joint (n=3,, t-test ,*p ≤ 0.05).

### Slide 2
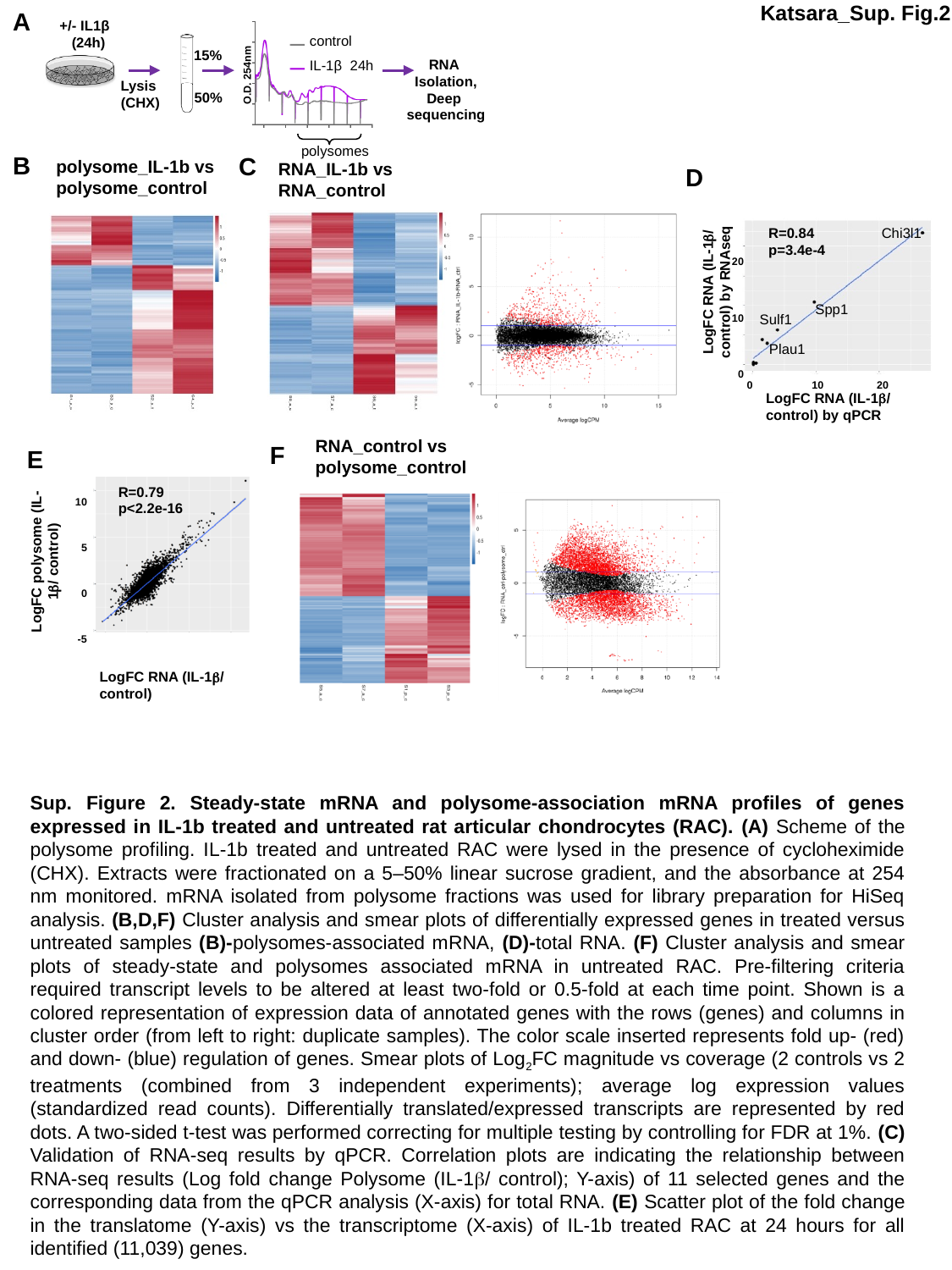

A
Katsara_Sup. Fig.2
+/- IL1β
 (24h)
control
IL-1β 24h
15%
RNA
Isolation,
Deep
sequencing
O.D. 254nm
Lysis
(CHX)
50%
polysomes
B
C
polysome_IL-1b vs
polysome_control
RNA_IL-1b vs
RNA_control
D
20
10
0
R=0.84
p=3.4e-4
Chi3l1
LogFC RNA (IL-1β/ control) by RNAseq
Spp1
Sulf1
Plau1
0 10 20
LogFC RNA (IL-1β/ control) by qPCR
RNA_control vs
polysome_control
F
E
10
5
0
-5
R=0.79
p<2.2e-16
LogFC polysome (IL-1β/ control)
LogFC RNA (IL-1β/ control)
Sup. Figure 2. Steady-state mRNA and polysome-association mRNA profiles of genes expressed in IL-1b treated and untreated rat articular chondrocytes (RAC). (A) Scheme of the polysome profiling. IL-1b treated and untreated RAC were lysed in the presence of cycloheximide (CHX). Extracts were fractionated on a 5–50% linear sucrose gradient, and the absorbance at 254 nm monitored. mRNA isolated from polysome fractions was used for library preparation for HiSeq analysis. (B,D,F) Cluster analysis and smear plots of differentially expressed genes in treated versus untreated samples (B)-polysomes-associated mRNA, (D)-total RNA. (F) Cluster analysis and smear plots of steady-state and polysomes associated mRNA in untreated RAC. Pre-ﬁltering criteria required transcript levels to be altered at least two-fold or 0.5-fold at each time point. Shown is a colored representation of expression data of annotated genes with the rows (genes) and columns in cluster order (from left to right: duplicate samples). The color scale inserted represents fold up- (red) and down- (blue) regulation of genes. Smear plots of Log_2_FC magnitude vs coverage (2 controls vs 2 treatments (combined from 3 independent experiments); average log expression values (standardized read counts). Differentially translated/expressed transcripts are represented by red dots. A two-sided t-test was performed correcting for multiple testing by controlling for FDR at 1%. (C) Validation of RNA-seq results by qPCR. Correlation plots are indicating the relationship between RNA-seq results (Log fold change Polysome (IL-1β/ control); Y-axis) of 11 selected genes and the corresponding data from the qPCR analysis (X-axis) for total RNA. (E) Scatter plot of the fold change in the translatome (Y-axis) vs the transcriptome (X-axis) of IL-1b treated RAC at 24 hours for all identified (11,039) genes.

### Slide 3
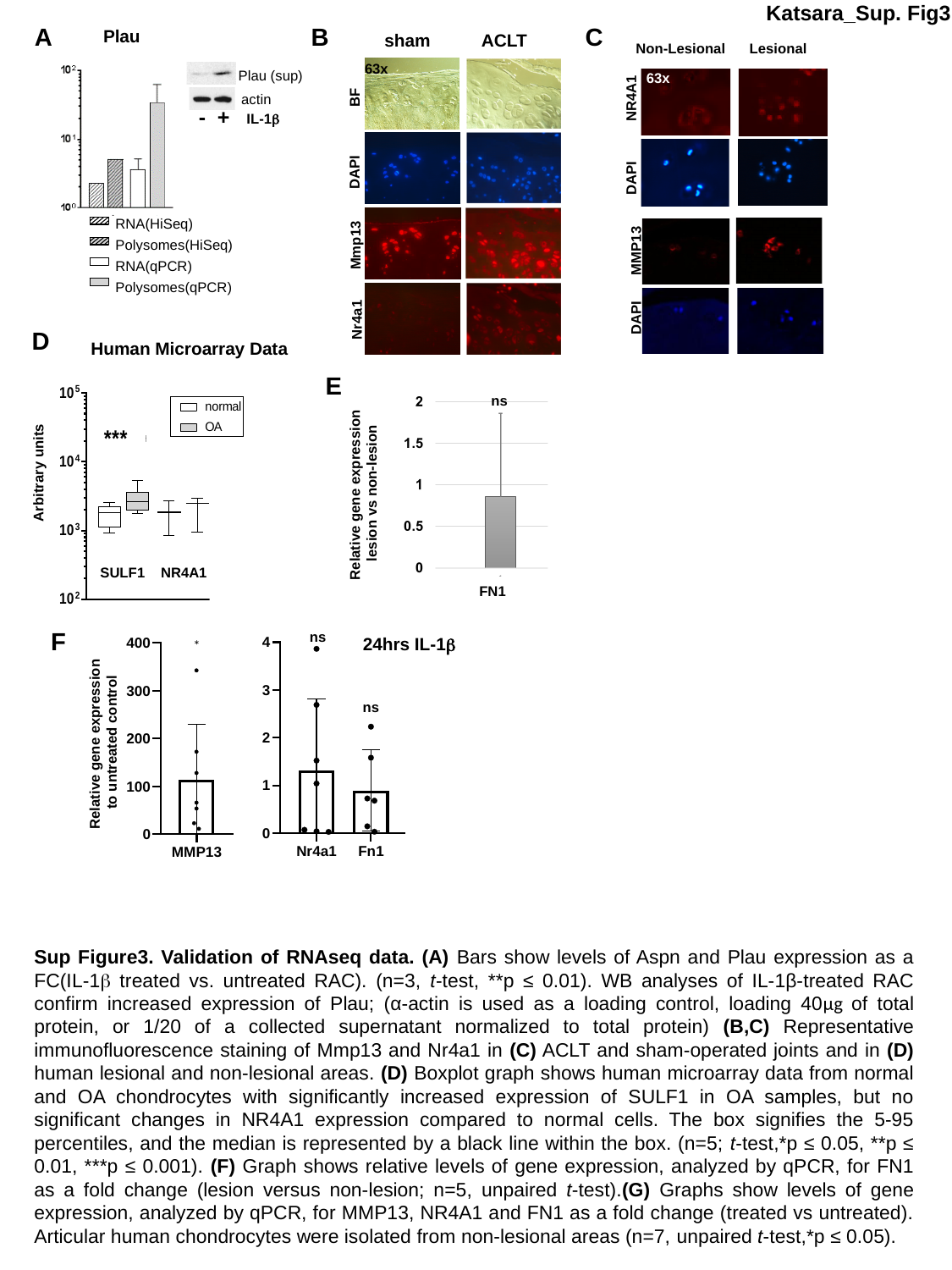

Katsara_Sup. Fig3
A
B
C
Plau
 Non-Lesional Lesional
 63x
NR4A1
DAPI
MMP13
DAPI
sham
ACLT
63x
BF
DAPI
Mmp13
Nr4a1
Plau (sup)
actin
IL-1β
- +
RNA(HiSeq)
Polysomes(HiSeq)
RNA(qPCR)
Polysomes(qPCR)
D
Human Microarray Data
E
ns
Arbitrary units
Relative gene expression
lesion vs non-lesion
SULF1 NR4A1
FN1
F
ns
*
24hrs IL-1β
ns
Relative gene expression
to untreated control
Sup Figure3. Validation of RNAseq data. (A) Bars show levels of Aspn and Plau expression as a FC(IL-1β treated vs. untreated RAC). (n=3, t-test, **p ≤ 0.01). WB analyses of IL-1β-treated RAC confirm increased expression of Plau; (α-actin is used as a loading control, loading 40µg of total protein, or 1/20 of a collected supernatant normalized to total protein) (B,C) Representative immunofluorescence staining of Mmp13 and Nr4a1 in (C) ACLT and sham-operated joints and in (D) human lesional and non-lesional areas. (D) Boxplot graph shows human microarray data from normal and OA chondrocytes with significantly increased expression of SULF1 in OA samples, but no significant changes in NR4A1 expression compared to normal cells. The box signifies the 5-95 percentiles, and the median is represented by a black line within the box. (n=5; t-test,*p ≤ 0.05, **p ≤ 0.01, ***p ≤ 0.001). (F) Graph shows relative levels of gene expression, analyzed by qPCR, for FN1 as a fold change (lesion versus non-lesion; n=5, unpaired t-test).(G) Graphs show levels of gene expression, analyzed by qPCR, for MMP13, NR4A1 and FN1 as a fold change (treated vs untreated). Articular human chondrocytes were isolated from non-lesional areas (n=7, unpaired t-test,*p ≤ 0.05).

### Slide 4
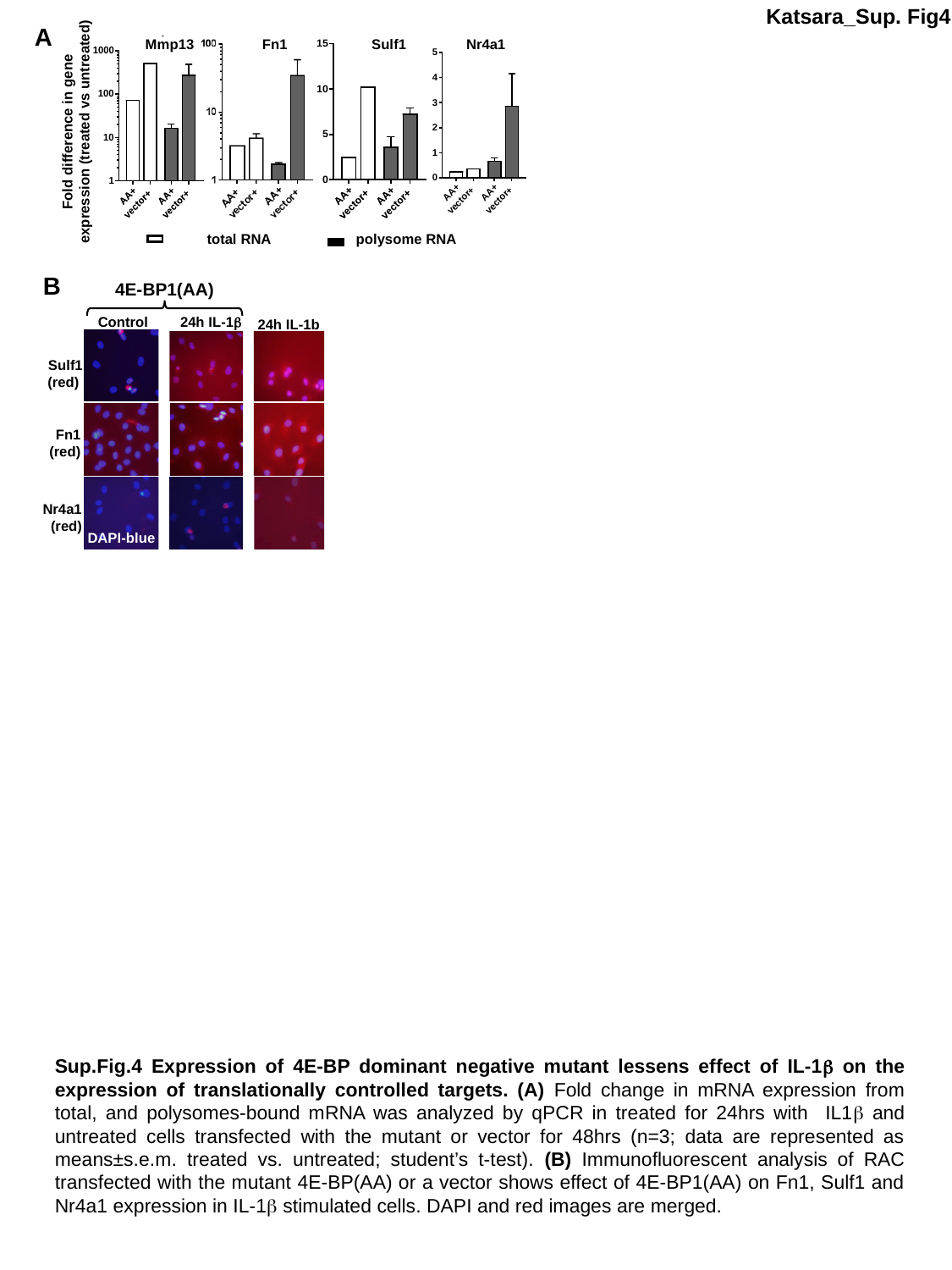

Mmp13 Fn1 Sulf1 Nr4a1
Fold difference in gene expression (treated vs untreated)
Katsara_Sup. Fig4
A
total RNA
polysome RNA
B
4E-BP1(AA)
Control 24h IL-1β
24h IL-1b
Sulf1
(red)
Fn1
(red)
Nr4a1
(red)
DAPI-blue
Sup.Fig.4 Expression of 4E-BP dominant negative mutant lessens effect of IL-1β on the expression of translationally controlled targets. (A) Fold change in mRNA expression from total, and polysomes-bound mRNA was analyzed by qPCR in treated for 24hrs with IL1β and untreated cells transfected with the mutant or vector for 48hrs (n=3; data are represented as means±s.e.m. treated vs. untreated; student’s t-test). (B) Immunofluorescent analysis of RAC transfected with the mutant 4E-BP(AA) or a vector shows effect of 4E-BP1(AA) on Fn1, Sulf1 and Nr4a1 expression in IL-1β stimulated cells. DAPI and red images are merged.

### Slide 5
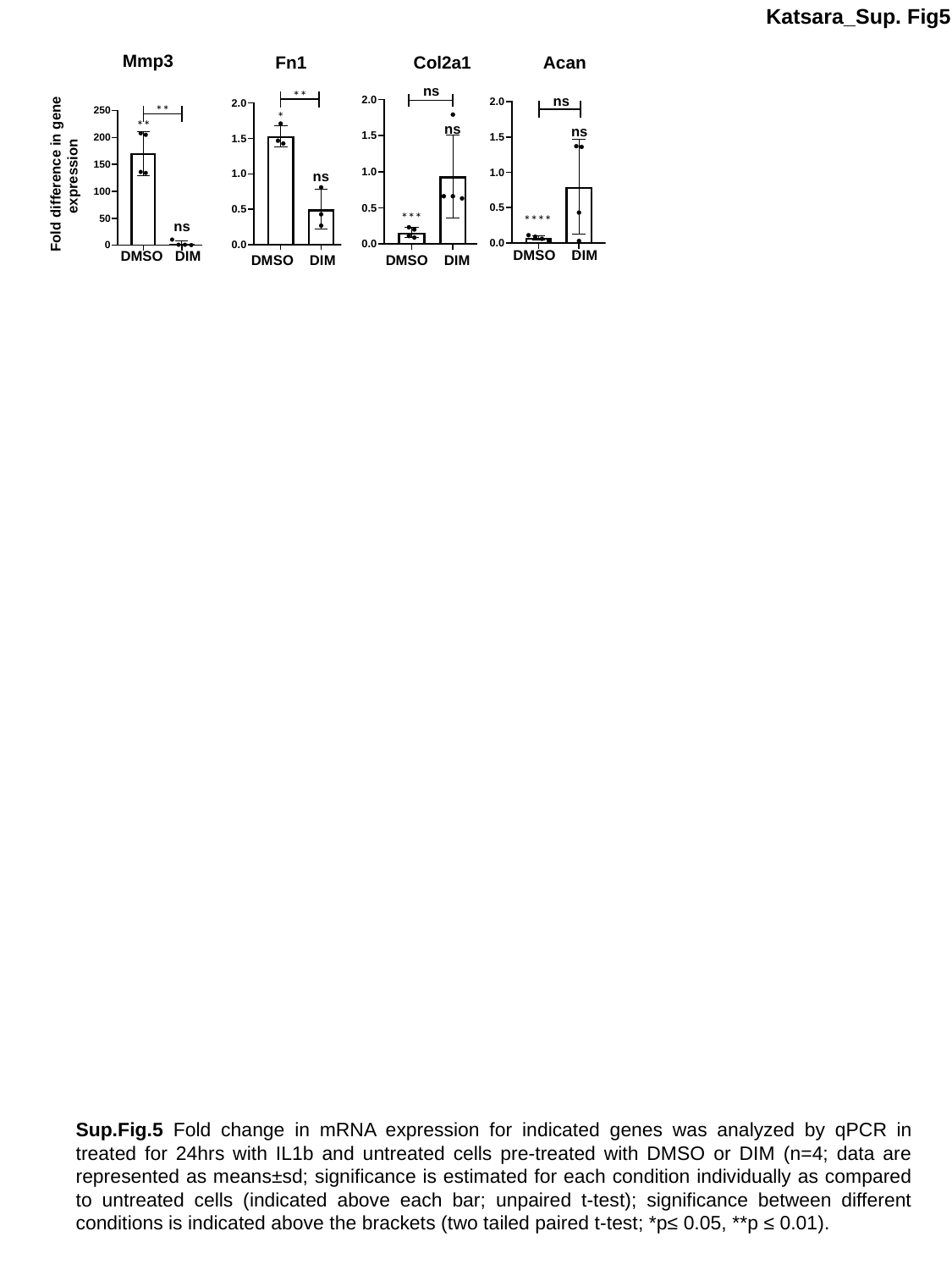

Katsara_Sup. Fig5
Mmp3
Fn1 Col2a1 Acan
ns
**
ns
*
ns
ns
ns
***
****
DMSO DIM
DMSO DIM
DMSO DIM
**
**
Fold difference in gene expression
ns
DMSO DIM
Sup.Fig.5 Fold change in mRNA expression for indicated genes was analyzed by qPCR in treated for 24hrs with IL1b and untreated cells pre-treated with DMSO or DIM (n=4; data are represented as means±sd; significance is estimated for each condition individually as compared to untreated cells (indicated above each bar; unpaired t-test); significance between different conditions is indicated above the brackets (two tailed paired t-test; *p≤ 0.05, **p ≤ 0.01).

### Slide 6
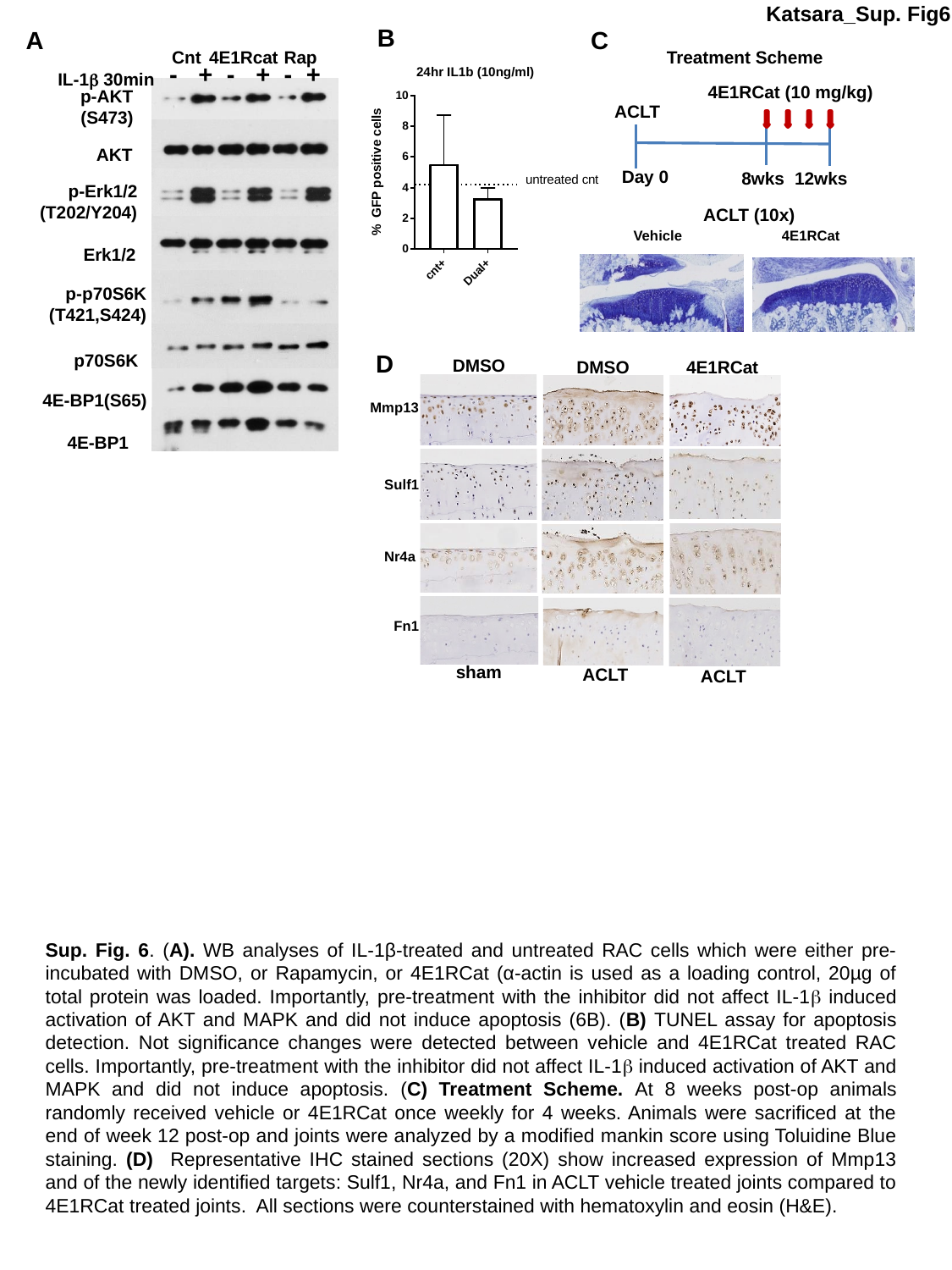

Katsara_Sup. Fig6
B
C
A
Cnt Rap
 4E1Rcat
Treatment Scheme
- + - + - +
IL-1β 30min
4E1RCat (10 mg/kg)
Day 0
8wks
12wks
p-AKT
 (S473)
ACLT
AKT
p-Erk1/2
(T202/Y204)
ACLT (10x)
Vehicle
4E1RCat
Erk1/2
p-p70S6K
(T421,S424)
D
p70S6K
DMSO
4E1RCat
DMSO
sham
ACLT
ACLT
4E-BP1(S65)
Mmp13
 4E-BP1
Sulf1
Nr4a
Fn1
Sup. Fig. 6. (A). WB analyses of IL-1β-treated and untreated RAC cells which were either pre-incubated with DMSO, or Rapamycin, or 4E1RCat (α-actin is used as a loading control, 20µg of total protein was loaded. Importantly, pre-treatment with the inhibitor did not affect IL-1β induced activation of AKT and MAPK and did not induce apoptosis (6B). (B) TUNEL assay for apoptosis detection. Not significance changes were detected between vehicle and 4E1RCat treated RAC cells. Importantly, pre-treatment with the inhibitor did not affect IL-1β induced activation of AKT and MAPK and did not induce apoptosis. (C) Treatment Scheme. At 8 weeks post-op animals randomly received vehicle or 4E1RCat once weekly for 4 weeks. Animals were sacrificed at the end of week 12 post-op and joints were analyzed by a modified mankin score using Toluidine Blue staining. (D) Representative IHC stained sections (20X) show increased expression of Mmp13 and of the newly identified targets: Sulf1, Nr4a, and Fn1 in ACLT vehicle treated joints compared to 4E1RCat treated joints. All sections were counterstained with hematoxylin and eosin (H&E).
